# Indication of a reduction in the cover of thin-leaved plants in Danish grasslands over an eight-year period

**DOI:** 10.1101/735472

**Authors:** Christian Damgaard

## Abstract

Across four grassland habitat types, the cover of thin-leaved plants was found to decrease significantly, but generally only limited trait selection was observed on leaf traits (SLA and LDMC) in a study of an extensive Danish grassland vegetation dataset from an eight-year period. The mostly negative result of this study may partly be due to the relatively conservative analysis, where the continuous plant trait variables are used for grouping plant species into functional types, which are then treated as dependent variables. This procedure is in contrast to most other analyses of trait selection, where it is the community weighted mean of the traits that is used as the dependent variable. However, it is not the traits, but rather individual plants that are sampled and, consequently, it is important to consider the sampling of species abundance in the statistical modelling of plant traits. This misapprehension has not received sufficient proper attention in the plant trait literature.

## Introduction

The observed covariation among plant traits and the environment has been the focus of plant ecology since its early days and has been demonstrated in numerus cases (e.g. Garnier *et al*., 2016). Furthermore, trait selection, i.e. a change in the distribution of plant traits in a specific area, has been observed in manipulated experiments (e.g. Laliberté *et al*., 2012, Pellissier *et al*., 2014) and during successional processes (e.g. Garnier *et al*., 2004, Garnier *et al*., 2007, Douma *et al*., 2012). Based on the above findings, it has been suggested to focus on the analysis of changes in mean plant traits rather than species abundance in order to understand community assembly rules and the causal mechanisms underlying species coexistence patterns (e.g. Stegen & Swenson, 2009, Shipley, 2010).

However, little work has been done to demonstrate trait selection in natural areas that are not subjected to large changes, such as successional processes. Conversely, in time-series analyses of plant species abundance in such natural areas it has frequently been demonstrated that species composition changes over time (e.g. Smart *et al*., 2006, Timmermann *et al*., 2015), but it has often been difficult to determine the causal mechanisms of the observed patterns of change in species abundance.

An often-studied suite of plant traits is the leaf economics spectrum, which is an important trade-off between plant resource acquisition and storage, where some plant species characterized by relatively thin leaves have a strategy of rapid carbon gain with rapid foliar turnover and fast growth. These thin-leaved species often have a competitive advantage in high-productive ecosystems with fertile soil. Whereas other plant species characterized by relatively thick leaves invest more resources into leaf construction and have a relatively slower foliar turnover and slow growth. These thick-leaved species often have a competitive advantage in low-productive ecosystems with infertile soil (e.g. Westoby *et al*., 2002, Craine, 2009, Reich, 2014).

In the past sixty years, there has been a dramatic anthropogenic increase in atmospheric nitrogen deposition, and significant effects on the plant vegetation have been demonstrated in some grassland habitats (Stevens *et al*., 2004, Damgaard *et al*., 2011), although nitrogen deposition recently has levelled off and even shown a decreasing tendency in Denmark (Ellermann *et al*., 2019). According to leaf economics spectrum theory, the increase in plant available nitrogen is expected to lead to selection in favor of thin-leaved fast growing plant species, although it is generally unknown how fast plant populations respond to environmental changes (Svenning & Sandel, 2013).

Usually, trait selection is investigated using community weighted mean trait values (CWMs) as the dependent variable; where the CWMs are calculated from a fixed species-trait matrix and the measured species abundances in a plot (e.g. Garnier *et al*., 2016). However, as pointed out by Clark (2016), it is not the CWMs that are sampled, but rather species abundances, and the stochastic properties of CWMs do not arise from variation in traits, but depend on the used sampling protocol to estimate species abundance. This fact that it is not the traits, but the individual plants that are sampled, is potentially critical for the statistical modelling of the covariance between traits and environment as well as trait selection (Clark, 2016), and this misapprehension has not received sufficient proper attention in the plant trait literature. One of the ways to solve the problem is to work with discrete plant functional types (PFTs) with known statistical sampling properties as the dependent variable rather than CWMs (Clark, 2016). However, depending on the studied problem it may not always be possible to define suitable PFTs, and then it is necessary to model the covariance between plant abundance and trait variation (Clark, 2016).

In this study, I will group grassland plant species according to leaf traits into PFTs and investigate whether the proportion among the abundance of the PFTs has changed over an eight-year period. The plant species will here be grouped with respect to two specific leaf traits, specific leaf area (SLA) and leaf dry matter content (LDMC), which are known to be important for determining the leaf economics spectrum and, consequently, the life history strategy of the plant species (e.g. Reich, 2014, Garnier *et al*., 2016).

## Materials and Methods

### Sampling design and plant cover data

Hierarchical time-series plant cover data in the eight-year period from 2007 to 2014 from 236 Danish grassland sites were used in the analysis. All sites included several grassland plots, with a total of 2946 plots that all were resampled three or more times with GPS-uncertainty (< 10 meters) over the sampling years. All plots at a site were either sampled or not sampled in a given year. Including resampling over the years, a total of 8859 vegetation plots were used in the analyses.

All plots were classified as belonging to one of four grassland habitat types: calcareous grasslands (or xeric sand calcareous grasslands, EU habitat type: 6120, 99 plots), dry grasslands (or semi-natural dry grasslands and scrubland facies on calcareous substrates, EU habitat type: 6210, 1155 plots), acid grasslands (or species-rich Nardus grasslands, EU habitat type: 6230, 1129 plots), and wet grasslands (or Molinia meadows on calcareous, peaty or clayey-siltladen soils, EU habitat type: 6410, 563 plots). The classification of the habitat type was performed according to the habitat classification system used for the European Habitat Directive (Nygaard *et al*., 2009, EU, 2013).

The plant cover, i.e. the relative projected area covered by a species, of all higher plants was measured by the pin-point method (Lindquist, 1931, Levy & Madden, 1933, Damgaard, 2009) using a square frame (50 cm X 50 cm) of 16 grid points that were equally spaced by 10 cm (Nielsen *et al*., 2012). One of the advantages of the pin-point method is that it is possible to aggregate taxa at the pin level. For example, if we want to aggregate the cover of two species, then we use the number of pins that hit either one or both of the species as an estimate of the aggregated cover of the two species.

The data are a subset of the ecological monitoring data collected in the Danish habitat surveillance program NOVANA (Nielsen *et al*., 2012, Nygaard *et al*., 2016).

### Leaf traits and species groups

The values of the leaf traits SLA and LDMC were found in the LEDA trait database (Kleyer *et al*., 2008), which is a homogenous trait database established from plant measurements in North Germany, which is geographically close to Denmark.

In order to group the higher plant species into PFTs, the trait values of the 542 grassland species where both leaf traits were available in the database were selected and analyzed in a principal component analysis (Fig. 1). The first principal component was used to group the species into four PFTs with similar mean abundance: 1: thin leaves (*PC*_1_ < 0), 2: intermediary leaves (0 ≤ *PC*_1_ < 1), 3: thick leaves (*PC*_1_ ≥ 1), 4: the rest, i.e. species where none or only one of the two leaf traits were available in the trait database.

**Fig. 1.**
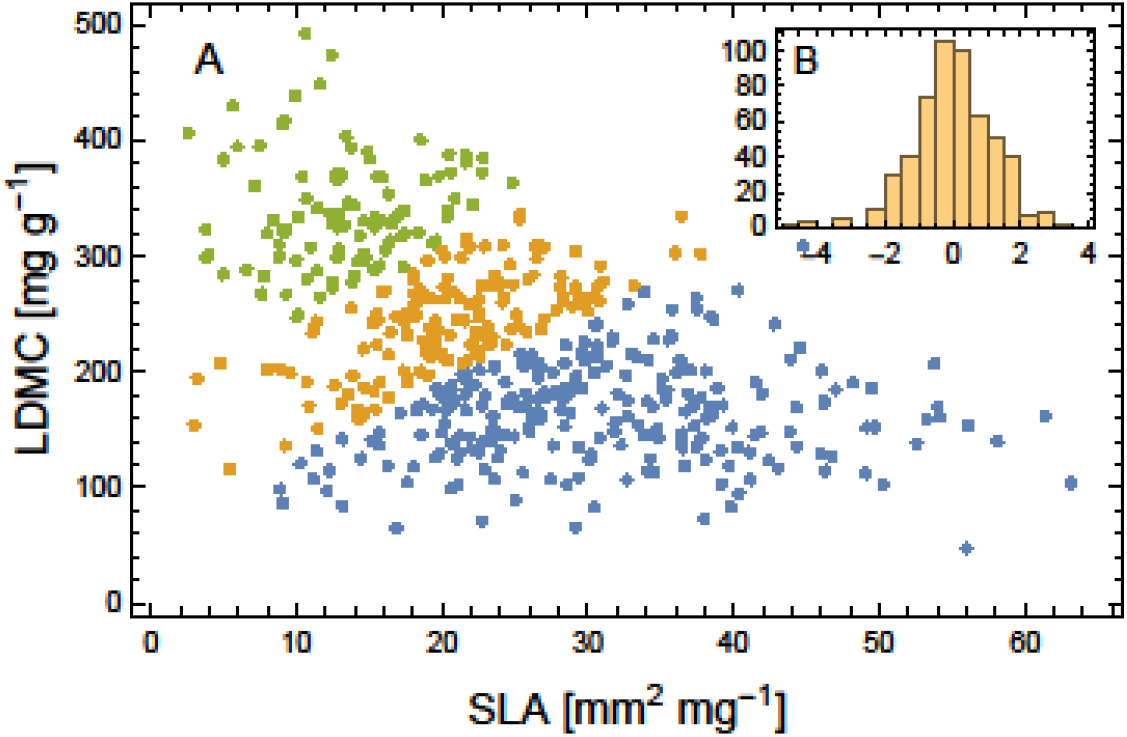
A: Leaf trait values of SLA and LDMC of the 542 grassland species, where both leaf traits were available in the database. PFTs: .thin leaves (blue), intermediary leaves (yellow) and thick leaves (green). B: histogram of the first principal component (*PC*_1_), which was used to group the plants into PFTs with similar overall mean abundance

The classification of each species is shown in the Electronic Supplement, and the mean site cover of the four different species groups is shown in Fig. 2.

**Fig. 2.**
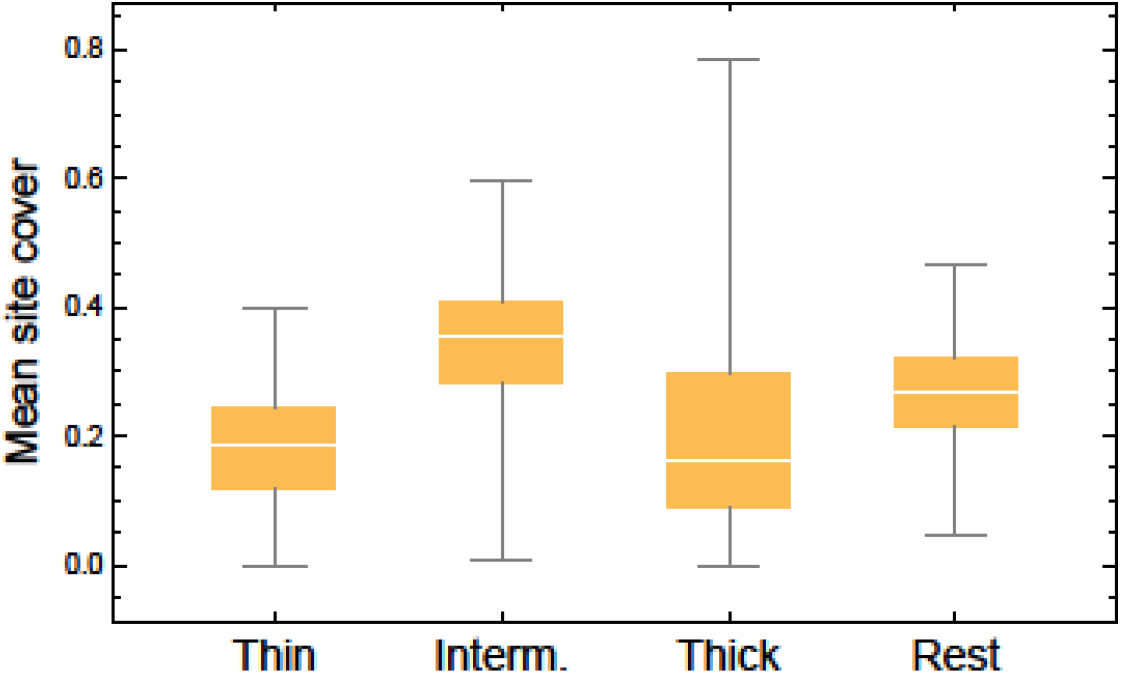
Distribution of the mean site cover of the four different species groups (PFTs).

**Fig. 3.**
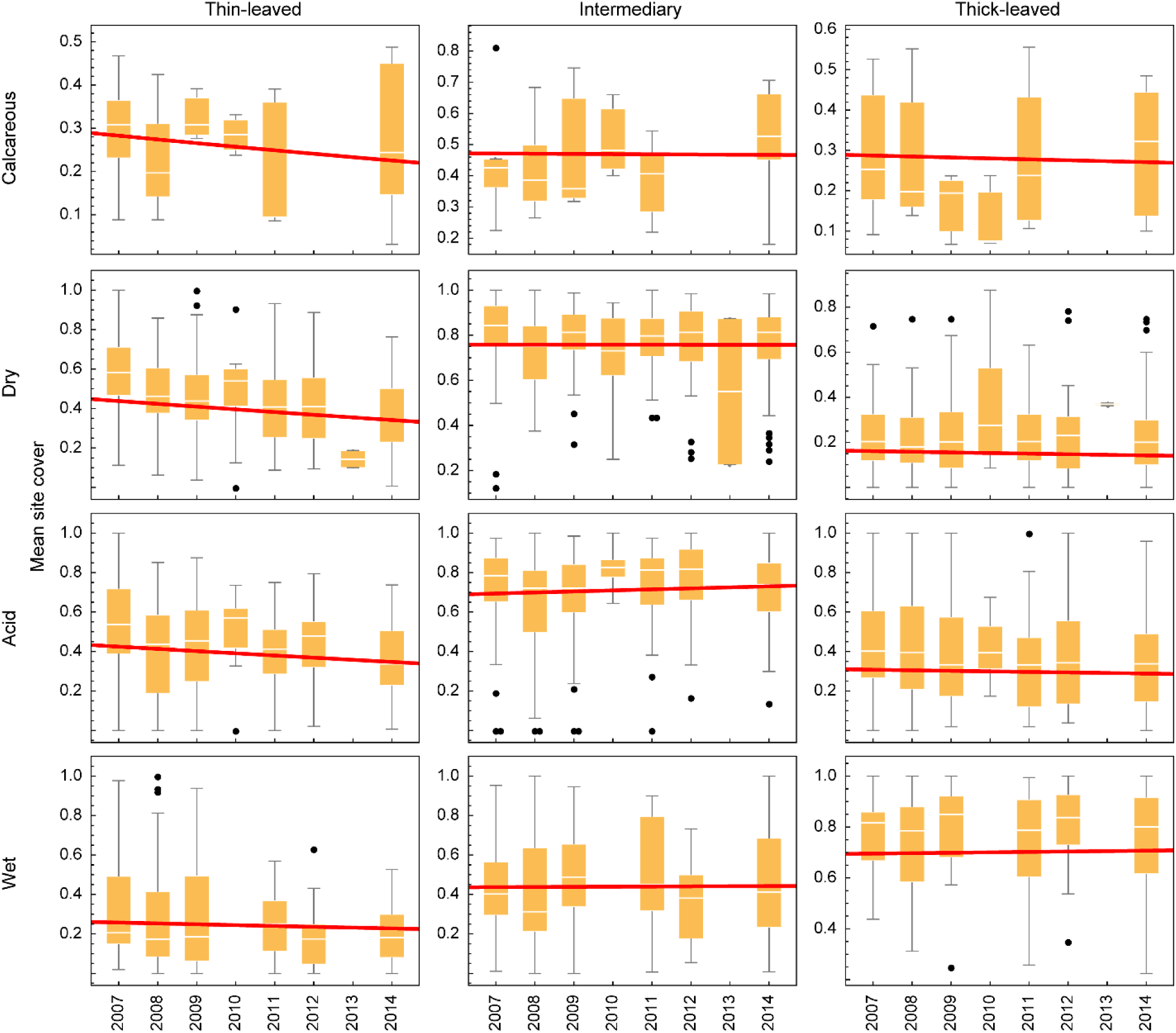
Distribution of the mean site cover at the four grassland habitat types over the years. The estimated slopes are illustrated with the red lines.

### Statistical model

The pin-point cover data are modelled using a reparametrized Dirichlet-multinomial distribution (Damgaard, 2015, Damgaard, 2018), where the mean plant cover of a species group *i* is assumed to be a linear model of year (*y*), habitat type (*j*) and site (*k*).

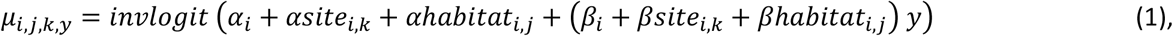

where site and habitat are modelled as random effects. Each random effect, consisting of *n* groups {1, …, *m*, …, *n*}, is modelled as *z_m_* = *σ ϵ_m_*, with a standard deviation (*σ* > 0) and *n* group regressors, *ϵ_m_*, that are assumed to have a strong prior distribution, *ϵ_m_*~*N*(0,1). For example, *βhabitat_i,j_* = *βhabitat_σ,i_ βhabitat_ϵ,i,j_*, where the parameter *βhabitat_σ,i_* is positive, and the *n_j_* parameters *βhabitat_ϵ,i,j_* are assumed to be standard normally distributed. Other location parameters had a weak normally distributed prior distribution. Standard deviation parameters and the degree of spatial aggregation had a uniform prior distribution in the ranges 0.01 to 10, and 0.01 to 0.9, respectively. In order to ensure that the sum of the mean cover parameters did not exceed one when the parameters in the linear model (1) were initialized to zero in the beginning of the MCMC sampling procedure, 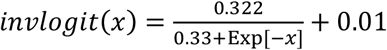 was used instead of transforming the linear model with the standard inverse logit function (Damgaard, 2018).

The model was fitted in STAN (Carpenter *et al*., 2017) using three chains of 100,000 iterations after a warmup of 300,000 iterations. The STAN model is included as an electronic supplement.

Statistical inferences were based on the marginal posterior distributions of parameters and compound parameters, i.e. their 2.5%, 50%, and 97.5% percentiles, and the proportion of distribution that is larger than zero, P(>0).

## Results

The fitted model had relatively many parameters, but, nevertheless, the MCMC iterations demonstrated good mixing properties and the marginal posterior distribution of the parameters had regular shapes (results not shown). Therefore, it was concluded that the model fitted the data adequately. The estimates of selected parameters and compound parameters are summarized in Tables 1, 2 and 3.

**Table 1.** Summary of the posterior distribution of the general parameters in model (1) for the different plant functional types (PFTs).

| Parameter | PFTs | Mean | SD | 2.5% | 50% | 97.5% | P(>0) |
| --- | --- | --- | --- | --- | --- | --- | --- |
| $\alpha$ | Thin-leaved | -0,558 | 0,474 | -1,449 | -0,572 | 0,626 | 0,068 |
| $\alpha$ | Intermediary | 0,255 | 0,386 | -0,660 | 0,264 | 1,034 | 0,834 |
| $\alpha$ | Thick-leaved | -0,710 | 0,712 | -2,151 | -0,700 | 0,802 | 0,111 |
| $\alpha_{site_{\sigma}}$ | Thin-leaved | 0,691 | 0,042 | 0,613 | 0,690 | 0,777 | - |
| $\alpha_{site_{\sigma}}$ | Intermediary | 0,538 | 0,033 | 0,477 | 0,536 | 0,608 | - |
| $\alpha_{site_{\sigma}}$ | Thick-leaved | 1,134 | 0,068 | 1,010 | 1,131 | 1,276 | - |
| $\alpha_{habitat_{\sigma}}$ | Thin-leaved | 0,744 | 0,653 | 0,225 | 0,537 | 2,934 | - |
| $\alpha_{habitat_{\sigma}}$ | Intermediary | 0,659 | 0,541 | 0,207 | 0,488 | 2,365 | - |
| $\alpha_{habitat_{\sigma}}$ | Thick-leaved | 1,325 | 0,923 | 0,439 | 1,041 | 3,983 | - |
| $\beta$ | Thin-leaved | -0,042 | 0,026 | -0,086 | -0,043 | 0,006 | **0,031** |
| $\beta$ | Intermediary | 0,006 | 0,025 | -0,036 | 0,006 | 0,054 | 0,688 |
| $\beta$ | Thick-leaved | -0,008 | 0,031 | -0,057 | -0,009 | 0,044 | 0,285 |
| $\beta_{site_{\sigma}}$ | Thin-leaved | 0,027 | 0,008 | 0,012 | 0,027 | 0,043 | - |
| $\beta_{site_{\sigma}}$ | Intermediary | 0,030 | 0,006 | 0,018 | 0,030 | 0,041 | - |
| $\beta_{site_{\sigma}}$ | Thick-leaved | 0,044 | 0,009 | 0,026 | 0,044 | 0,063 | - |
| $\beta_{habitat_{\sigma}}$ | Thin-leaved | 0,036 | 0,043 | 0,011 | 0,025 | 0,134 | - |
| $\beta_{habitat_{\sigma}}$ | Intermediary | 0,032 | 0,036 | 0,010 | 0,021 | 0,119 | - |
| $\beta_{habitat_{\sigma}}$ | Thick-leaved | 0,036 | 0,048 | 0,011 | 0,023 | 0,134 | - |

**Table 2.** Summary of the posterior distribution of the change in cover (*β_i_* + *βhabitat_σ,i_ βhabitat_ϵ,i,j_*) for the different plant functional types (PFTs) in the different grassland habitat types

| Habitat type | PFTs | Mean | SD | 2.5% | 50% | 97.5% | P(>0) |
| --- | --- | --- | --- | --- | --- | --- | --- |
| calcareous grasslands | Thin-leaved | -0,046 | 0,065 | -0,168 | -0,046 | 0,081 | 0,175 |
| calcareous grasslands | Intermediary | -0,012 | 0,079 | -0,160 | -0,011 | 0,132 | 0,422 |
| calcareous grasslands | Thick-leaved | -0,012 | 0,081 | -0,170 | -0,012 | 0,146 | 0,409 |
| dry grasslands | Thin-leaved | -0,060 | 0,058 | -0,178 | -0,059 | 0,051 | 0,090 |
| dry grasslands | Intermediary | -0,002 | 0,075 | -0,141 | -0,002 | 0,135 | 0,486 |
| dry grasslands | Thick-leaved | -0,022 | 0,078 | -0,174 | -0,021 | 0,128 | 0,324 |
| acid grasslands | Thin-leaved | -0,055 | 0,058 | -0,172 | -0,055 | 0,057 | 0,103 |
| acid grasslands | Intermediary | 0,018 | 0,074 | -0,122 | 0,017 | 0,157 | 0,654 |
| acid grasslands | Thick-leaved | -0,012 | 0,078 | -0,160 | -0,012 | 0,139 | 0,395 |
| wet grasslands | Thin-leaved | -0,033 | 0,060 | -0,148 | -0,034 | 0,085 | 0,202 |
| wet grasslands | Intermediary | -0,009 | 0,075 | -0,152 | -0,007 | 0,126 | 0,441 |
| wet grasslands | Thick-leaved | 0,010 | 0,079 | -0,140 | 0,007 | 0,167 | 0,566 |

**Table 3.** Summary of the posterior distribution of the degree of spatial aggregation in the different grassland habitat types.

| Habitat type | Mean | SD | 2.5% | 50% | 97.5% |
| --- | --- | --- | --- | --- | --- |
| calcareous grasslands | 0,122 | 0,007 | 0,110 | 0,122 | 0,136 |
| dry grasslands | 0,103 | 0,002 | 0,100 | 0,103 | 0,107 |
| acid grasslands | 0,110 | 0,002 | 0,106 | 0,110 | 0,114 |
| wet grasslands | 0,124 | 0,003 | 0,118 | 0,124 | 0,131 |

Across the four grassland habitat types, the cover of thin-leaved plants decreased significantly during the eight-year period (Table 1: *β* – thin-leaved, *P* = 0.031). However, within the four grassland habitat types there were no significant changes in the cover of any of the PFTs (Table 2); the strongest statistical signal was a negative trend for thin-leaved plant species in dry grasslands (Table 2: thin-leaved, *P* = 0.090).

The estimated standard deviations of the yearly change among sites were approximately of the same size as the standard deviations among habitats (Table 1), which indicates that there is a sizeable variation in the yearly change among sites of the same grassland habitat type.

The standard deviations of the posterior distributions of the slope parameters that measure the yearly change in the cover of the PFTs were approximately 0.03 on the logit scale (Table 1). This width of the posterior distribution indicates that the statistical power of the analysis is moderate, and in connection with the mostly non-significant changes in cover, it is concluded that trait selection for the studied leaf traits in Danish grasslands over the eight-year period generally is limited.

The spatial aggregation of the FPTs were relatively benign and did not differ significantly among the grassland habitat types (Table 3).

## Discussion

Across the four grassland habitat types, the cover of thin-leaved plants was found to decrease significantly in this study of an extensive eight-year Danish grassland vegetation dataset, and the hypothesis that the increase in plant available nitrogen the past sixty years should have selected for thin-leaved fast growing plant species cannot be corroborated. This result may indicate that the observed decrease in nitrogen deposition over the last years is beginning to reverse earlier demonstrated effects of elevated nitrogen deposition on plant traits, but there are other possible explanations such as changed management, increase in soil moisture, and reduced grazing pressures on the studied grasslands (Cruz *et al*., 2010, Zheng *et al*., 2015). The result of the present study is also in contrast to the results reported by Timmermann et al. (2015), who in a shorter time-series observed that the winning plant species, i.e. the species that had an increasing cover, were relatively thin-leaved (had a relatively small SLA). Generally, more times-series studies of trait selection in natural areas that are not subjected to large changes, such as successional processes, are needed in order to validate the predictions made in space-for-time studies of plant traits (Damgaard, 2019).

The chosen method of analysis, where the continuous variables SLA and LDMC are used for grouping plant species into PFTs and the aggregated cover of the PTFs are then treated as the dependent variable instead of treating the CWMs of SLA and LDMC as the dependent variable, is relatively conservative. However, as mentioned in the introduction it is not the traits, but the individual plants that are sampled, and it is important to consider the sampling of species abundance in the statistical modelling of plant traits (Clark, 2016).

An alternative method of analyzing plant trait selection is to correlate trait values with estimated demographic parameters or change in abundance for individual species (Timmermann *et al*., 2015, Garnier *et al*., 2018, Herben *et al*., 2019). However, it may be a problem that typically only point estimates of the plant ecological success are used without considering the covariance between plant abundance and trait variation (Clark, 2016). Furthermore, since many plant traits are correlated (e.g. Reich, 2014) it is problematic to infer from observed correlations to causal mechanisms. Especially since the studied plant traits often are the traits that are most readily available in trait databases.

In this study, plant abundance is measured by plant cover using the pin-point method, which readily allows the aggregation of single cover into the cover of PFTs at the pin level. As demonstrated elsewhere, it is important to take the spatial aggregation of plant species into account when modelling plant cover (Damgaard, 2012, Damgaard, 2013, Damgaard & Irvine, 2019), and these results have been generalized into a multi-species setting, where it is recommended to analyze multispecies or PFTs pin-point cover data in a reparametrized Dirichlet-multinomial distribution (Damgaard, 2015, Damgaard, 2018). A further advantage of modelling trait selection by the change in the abundance of PFTs rather than the change in the continuous CWMs is that the class of missing values, i.e. the plant species with no or incomplete information on the trait values, is clearly defined (in this study as the rest group). Additionally, more complicated selection models than directional selection may be readily tested, e.g. stabilizing selection (the intermediary PFT is positively selected) and disruptive selection (the intermediary PFT is negatively selected).

The studied vegetation plots were not real permanent plots, but were only resampled with GPS-uncertainty. However, if we had access to permanent plot time-series cover data it would have been interesting to explore more sophisticated selection models, e.g. whether the observed trait selection could be partitioned into direct selection and selection that is mediated by interspecific interactions (Damgaard, 2016, Pedersen *et al*., 2019).

## Supporting information

Species list (Excel file)

STAN model (text file)

## Electronic Supplements

A: Species list (Excel file)

B: STAN model (text file)

